# Replication coordination marks the domestication of large extrachromosomal replicons in bacteria

**DOI:** 10.1101/2025.03.15.643453

**Authors:** Jakub Czarnecki, Morgan Lamberioux, Ole Skovgaard, Amaury Bignaud, Najwa Taib, Théophile Niault, Pascale Bourhy, Julia Bos, Elvira Krakowska, Dariusz Bartosik, Romain Koszul, Martial Marbouty, Didier Mazel, Marie-Eve Val

## Abstract

Bacterial genomes often include extrachromosomal replicons (ERs), ranging from small plasmids to nearly chromosome-sized elements, that foster genome plasticity and adaptation. Despite their prevalence, the mechanisms underlying ER domestication and their long-term adaptation within bacterial hosts remain largely unexplored. By analyzing over 40,000 complete bacterial genomes, we identified two main ER categories: small ERs with diverse GC content and large ERs (≥10% the size of the main chromosome) that closely match the GC content of the chromosome. Across multiple phyla, marker frequency analyses showed that large ERs maintain a 1:1 copy number with the chromosome. Another key finding of this study is that large ERs terminate replication in synchrony with the chromosome. Hi-C contact maps revealed consistent *ori*-*ori* interactions between chromosomes and ERs. In large ERs, inter-replichore and *ter*-*ter* interactions, along with the recruitment of key chromosomal segregation motifs, suggest the co-option of chromosome-associated replication and segregation machineries. Together, our findings indicate that as ERs become larger, they become increasingly reliant on chromosome-driven processes for stable inheritance, potentially explaining why they do not exceed the size of the chromosome.

## INTRODUCTION

The bacterial genome undergoes dynamic structural and functional changes throughout the cell cycle, following a repeatable sequence of events governed by its maintenance systems. At the core of these events is the fate of the main chromosome, an essential replicon conserved across bacterial evolutionary lineages ^1^. Alongside the chromosome, many bacteria harbor extrachromosomal replicons (ERs), which exhibit remarkable diversity in size, GC content, copy number, evolutionary conservation, and their interactions with the host cell ^2^.

### Plasmids and their evolution into larger replicons

Small ERs, commonly known as plasmids, are the most widespread and variable ERs among bacteria. Typically non-essential, plasmids can be lost by their host without affecting viability. They are often viewed as molecular parasites due to their potential to impose an energetic burden on their hosts ^3–5^. However, plasmids also play a significant role in host fitness by carrying adaptive genes, such as those conferring antibiotic resistance or niche-specific traits, and by increasing the genomic flexibility of their hosts without disturbing the synteny ^5,6^. To ensure inheritance, plasmids rely on a high copy number, which increases the likelihood of being present in both daughter cells after division, and/or encoding their own partitioning systems ^7,8^. Plasmids are often mobile, capable of horizontal transfer via conjugation ^9^ within bacterial communities, a mechanism that further enhances their stability and persistence within populations ^10^.

Through genome shuffling and DNA acquisition, plasmids can evolve into larger structures, giving rise to megaplasmids and chromids ^11–13^. This process involves acquiring additional DNA while simultaneously lowering copy numbers to reduce the metabolic burden on the host cell. To maintain stability, both megaplasmids and chromids can rely on host genome maintenance systems. The primary distinction between these replicons lies in their interaction with the host.

Megaplasmids, although not essential, often carry critical genetic information that supports survival in the host’s specific ecological niche ^13^. Retaining their mobility, megaplasmids can spread horizontally through conjugation, which accounts for their lack of conservation even within the same species ^11,12^. In contrast, chromids carry core essential genes ^11,12,14,15^. Unlike megaplasmids, chromids lose their mobility, and fully rely on the vertical transmission ^9^. This integration confers chromosome-like stability, making chromids highly conserved within certain evolutionary lineages, often at the genus level ^11,12^.

### Challenges in ER classification

The classification of bacterial ERs remains complex despite established frameworks, such as those proposed by Harrison et al. (2010), DiCenzo and Finan (2017), and Hall et al. (2021) ^11–13^. Chromosomes are consistently identified as the largest replicons, characterized by a DnaA-dependent replication system ^11,16^. ERs (including plasmids, megaplasmids, and chromids) are typically defined by features such as size, GC content, mobility, and the absence or presence of essential core genes ^11–13^. However, these frameworks are often limited by arbitrary thresholds (e.g., a 350 kb size cutoff between plasmids and megaplasmids) ^12,13^ and difficulties in defining the category of essentiality. This challenge arises because the essentiality of a replicon is often dependent on specific growth conditions, with some replicons encoding highly conserved functions that are critical in a given environment but not universally essential ^17,18^.

Despite the existence of these classifications, they are not implemented when bacterial replicons are deposited in the RefSeq database. Instead, replicon names are assigned by the researchers depositing the genomes without standardized guidelines. Consequently, most ERs are broadly categorized as plasmids, while only a few are labeled as megaplasmids. Additionally, some particularly large ERs are designated as secondary chromosomes, even when evidence suggests they are more closely related to plasmids ^11^. This inconsistency stems from the fact that current definitions of plasmids, megaplasmids, and chromids are challenging to apply in practice.

The number of complete bacterial genomes available has grown significantly since the development of the abovementioned ER classification frameworks. This increase in genomic data provides an important opportunity to revisit these classifications and work toward a simpler, more universal system.

### Studies on behavior of large ERs

NGS-based techniques, including Marker Frequency Analysis (MFA) ^19^ and 3C/Hi-C ^20^, have advanced our understanding of large ER maintenance and behavior. Studies on model organisms, such as *Vibrio cholerae* (and other Vibrionaceae), *Burkholderia cepacia*, *Agrobacterium tumefaciens*, and *Pseudoalteromonas* spp., have revealed diverse strategies for ER integration into genome maintenance systems ^21–28^. MFA consistently shows that large ERs synchronize their replication termination with the main chromosome, indicating coordinated replication ^21,25,26^. Complementary 3C/Hi-C analyses reveal physical interactions between replicons. In *V. cholerae*, the chromid origin of replication (*ori2*) interacts with *crtS*, a chromosomal sequence whose replication triggers chromid replication initiation, ensuring termination synchrony ^21,29^. Additionally, weak *ter*-*ter* interactions have been observed between the chromosome and chromid ^21,30^. In *A. tumefaciens*, Hi-C data highlight interactions among all replication origins, chromosome, chromid, megaplasmid, and plasmid ^24^. These findings underscore the diversity of strategies bacteria use to integrate large ERs, though the extent of these mechanisms across taxa remains unclear.

To address significant gaps in our knowledge of large ERs, we conducted a comprehensive investigation aimed at refining their classification, mapping their distribution, and uncovering the mechanisms that govern their maintenance and interactions with chromosomes. **(1) <u>ER classification and distribution across bacteria</u>:** We analyzed all available complete bacterial genomes from RefSeq to develop a universal framework for classifying extrachromosomal replicons (ERs). By focusing on features such as size and GC content, we aimed to simplify classification and examine the distribution and prevalence of ER types across bacterial taxa. **(2) <u>Replication</u> <u>coordination</u>:** Using MFA, we investigated the replication dynamics of large ERs and observed that replication termination synchronization, is conserved across diverse bacterial species. **<u>(3) Chromosome-ER interactions</u>:** Hi-C analysis allowed us to explore physical interactions between chromosomes and ERs. By building on insights from model organisms such as *V. cholerae* and *A. tumefaciens*, we showed that similar patterns of replication and segregation are conserved across bacterial lineages, shedding light on the integration of large ERs into genome maintenance systems.

This study combines large-scale bioinformatics with targeted experimental approaches to provide new insights into the structure and function of bacterial genomes. Our findings establish a foundation for future research into multipartite bacterial genomes and the domestication of extrachromosomal elements.

## RESULTS

### ERs can be divided in two distinct groups

We analyzed a database of over 40,000 complete bacterial genomes from RefSeq to develop methods for classifying bacterial replicons and to provide a comprehensive overview of ER abundance across bacterial taxa.

### Chromosomes vs. ERs

First, we evaluated whether chromosomes and ERs form distinct groups in two-dimensional analysis based on their GC content and size. Here we define chromosome as the largest replicon in the genome, which is a convenient proxy supported by the observation that replicons with conserved chromosome-type DnaA-based replication systems are consistently the largest in bacterial genomes ^11,12^. All other replicons were classified as ERs. Our analysis, stratified by phyla demonstrates that chromosomes and ERs form well-separated clusters in most cases (Fig. 1a). Overlap between these two groups is observed primarily in Pseudomonadota, a phylum known to be especially rich in large ERs.

**Figure 1:**
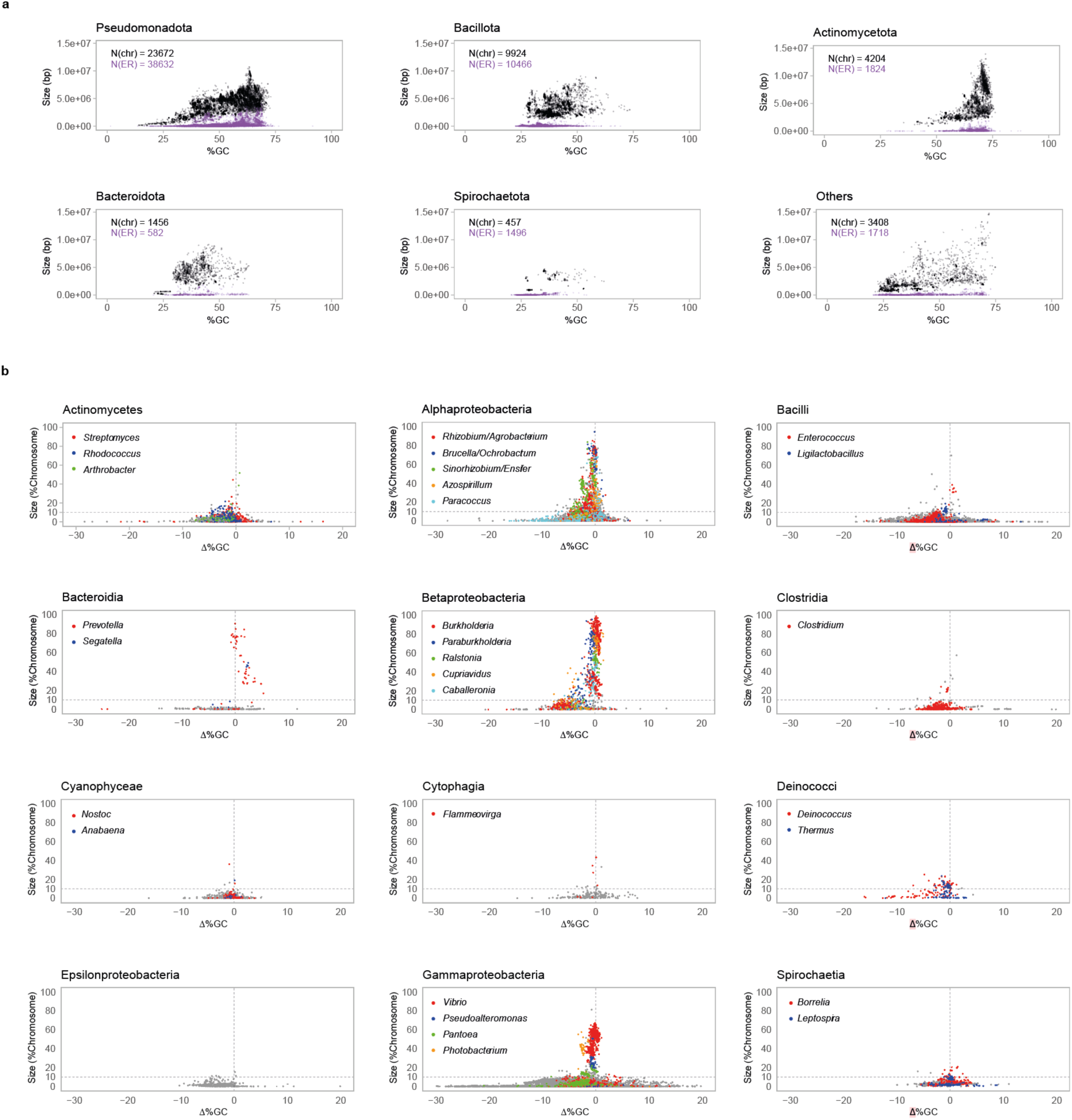
**a. Distribution of chromosomes and extrachromosomal replicons (ERs) by phylum based on GC content and size.** Chromosomes (black) and extrachromosomal replicons (ERs; purple) from the five phyla with the highest number of ERs in the RefSeq database, along with data from other phyla, are analyzed. Replicons are plotted by size (bp) and GC content (%), using a continuous scale. The total numbers of chromosomes (N(chr)) and ERs (N(ER)) are indicated on each plot. The number of chromosomes corresponds to the number of genomes analyzed per plot, as chromosomes were defined as the largest replicons in each genome, while ERs were categorized as all other replicons within the same genome. **b. Classification of Extrachromosomal Replicons by size and GC content across taxonomic classes.** ERs from bacterial taxonomic classes with more than 100 ERs in our dataset are plotted based on their size, expressed as a percentage of the chromosome (%Chromosome), and the difference in GC content compared to the chromosome of the same cell (Δ%GC). For each class, ERs from up to five genera with the highest number of large ERs (defined as those exceeding 10% of the chromosome size) are highlighted. Dashed lines indicate the 10% Chr threshold, which serves as our proposed boundary between small and large ERs, and Δ%GC = 0, where ERs share the same GC content as their corresponding chromosome. **Extended Legend Fig. 1:** All genera with at least eight genomes, where ≥50% contain large ERs are listed below, and also other important examples are included. For each genus, we report the number of genomes with large ERs relative to the total number of genomes from that genus in our dataset (indicated as XX/XX): **Gammaproteobacteria:** Large ER clusters correspond to secondary chromosomes in *Vibrio* (601/604), *Photobacterium* (41/41), and *Pseudoalteromonas* (75/82), and megaplasmids in *Pantoea* (70/107). All *Vibrionaceae* genera harbor conserved Chr2, including *Allivibrio* (5/5), *Grimontia* (6/6), *Paraphotobacterium* (1/1), *Salinivibrio* (7/7), and *Thaumasiovibrio* (1/1). **Betaproteobacteria:** Large ERs in *Burkholderia* (462/467) form two clusters corresponding to Chr2 and Chr3. Other major clusters include Chr2 of *Paraburkholderia* (53/54), *Ralstonia* (74/74), *Cupriavidus* (42/43), and *Caballeronia* (12/12). *Mycetohabitans* (9/9) also shows high prevalence of large ERs. **Alphaproteobacteria:** *Sinorhizobium/Ensifer* (85/85) has two clusters of large ERs: pSymB-like (Δ%GC ≈ 0) and pSymA-like (Δ%GC ≈ -2.5). Other large ERs represent Chr2 in *Brucella/Ochrobactrum* (220/222) and megaplasmids in *Azospirillum* (19/19). Additional conserved large ERs appear in *Rhizobium/Agrobacterium* (261/274), and *Paracoccus* (27/52). In this class, many more genera meet the eight genomes and ≥50% threshold, including *Cereibacter* (9/9), *Tistrella* (17/17), *Ruegeria* (8/9), *Shinella* (10/13), *Novosphingobium* (16/22), and *Roseomonas* (8/16). **Spirochaetia:** *Leptospira* has two large ER groups: smaller plasmids and larger chromosome 2, often just below the 10% Chr threshold. *Borrelia* also carries large ERs but lacks a distinct cluster. **Bacilli & Clostridia:** Clusters of Large ERs (Δ%GC ≈ 0) in *Enterococcus* (17/697) and *Clostridium* (20/358) suggest rare domesticated ERs in Gram-positive bacteria. The low prevalence of large ERs in these genera suggests that their conservation might be species-specific. For instance, in Clostridium, large ERs are mainly found in *Clostridium butyricum*. **Bacteroidia & Deinococcus-Thermus:** Large ERs are frequently found in Prevotella (47/65) and Segatella (4/8) within Bacteroidia, and in Deinococcus (31/39) and Thermus (30/43) within Deinococcus-Thermus. **Actinomycetes:** Few *Streptomyces* (30/914) carry large ERs, and also in *Arthrobacter* (4/73) and *Rhodococcus* (18/136). **Cyanophyceae:** Rare large ERs in *Nostoc* (2/22) and *Anabaena* (1/4). **Cytophagia:** Poorly sampled; *Flammeovirga* (4/4) is the only genus with conserved large ERs. **Epsilonbacteria:** Large ERs are nearly absent.

### Delineation of two ER groups

Distinguishing ERs clusters by size and GC content at the phylum level is nearly impossible (Fig. 1a, Supplementary Fig. 1). To address this, we excluded main chromosomes and analyzed only ERs, exploring alternative ways to express their size and GC content. For size, we measured it in absolute numbers (bp), as a percentage of the whole genome and as a percentage of the chromosome. Expressing replicon size as a percentage of the whole genome was proposed by Hall *et al.* ^13^, based on the rationale that size-based ER classification should account for the variability in genome sizes across bacteria. We extended this approach by using chromosome size as a reference instead of genome size, reasoning that the chromosome is the most conserved part of the genome and provides a more robust and stable measure for comparisons. While ERs can contribute significantly to the total genome size, their profiles often vary between strains of the same species. For GC content, we expressed it as a total value and as the difference with that of the chromosome (Δ%GC). The rationale for this approach is that bacterial chromosomes differ widely in GC content, but ERs undergoing co-evolution with their host chromosome tend to have GC content closer to the chromosome. Calculating Δ%GC removes bias caused by differences in the baseline GC content of various bacteria and helps highlight co-evolutionary patterns of ER domestication across different species.

Using these three size measures (absolute size, percentage of genome, and percentage of chromosome) and two GC content measures (total GC content and Δ%GC), we generated six comparative plots (Supplementary Fig. 2). These plots were prepared across families, with ERs color-coded based on their current classification in the RefSeq database. As a validation of our approach, we examined the Burkholderiaceae family, a group of bacteria with a well-characterized genome structure, including two conserved secondary chromosomes (Chr2 and Chr3), as well as numerous plasmids^31^. Our method accurately classified these replicons (Supplementary Fig. 2 - Burkholderiaceae). Specifically, using Δ%GC as a GC content measure, rather than %GC, allowed a better distinction of Burkholderiaceae Chr2 and Chr3 into well-defined clusters in the lower set of plots (Δ%GC). While the differences between size measures were less pronounced, the best resolution between small plasmids (orange) and secondary chromosomes (blue) was achieved when size was expressed as a percentage of the chromosome (%Chr). This pattern was consistent across other families (Supplementary Fig. 2 – *Rhizobiaceae*, *Vibrionaceae*). Based on these observations, we selected size expressed as %Chr and GC content expressed as Δ%GC for our subsequent analyses.

We applied this approach to plot ERs from 12 taxonomic classes with more than 100 ERs in the RefSeq database (Fig. 1b), as well as families with more than 100 ERs to provide a more detailed taxonomic resolution (Supplementary Fig. 3). In most classes, we observed a bell-shaped distribution of ERs, with Δ%GC decreasing as ER size increases. The distribution peaks around 10 %Chr, slightly below chromosomal %GC. However, many classes also include larger ERs (>10 %Chr) with Δ%GC values close to zero, which are particularly abundant in Alpha-, Beta-, and Gammaproteobacteria (Fig. 1b). This analysis allowed us to identify two main ER categories: (1) <u>Small ERs</u>: defined as those with sizes below 10 %Chr and diverse Δ%GC values and (2) <u>Large ERs</u>: defined as those with sizes equal to or greater than 10 %Chr and Δ%GC values near zero.

### Distribution and conservation of large ERs

In most classes, ER groups are easily distinguishable in the plots. However, distinguishing them is more challenging in Alphaproteobacteria due to the presence of many replicons occupying the intermediate space between these two groups. To refine our analysis, we identified genera with a high percentage of genomes carrying large ERs (>10% Chr), prioritizing well-sampled genera for robust observations. We selected up to five genera per class (Fig. 1b), focusing on those where large ERs are conserved. Albeit some classes lacked such genera, representative examples were still included. In many cases, large ERs from the same genus clustered together, suggesting the presence of conserved large ERs within these genera. In contrast, more scattered distributions indicate either less conserved ERs or genera with fewer representatives in our dataset. In addition to the genera highlighted in Fig. 1b, we listed all genera with at least eight genomes, where ≥50% contain large ERs (Fig. 1 Extended Legend, Supplementary Data 2). Our analysis shows that conserved large ERs are mainly found in certain proteobacterial genera, while their distribution in other classes is more sporadic. The 10% Chr threshold may require refinement for specific taxa where conserved ERs exist but fall just below the large ER threshold (e.g. *Leptospira*). While most large ERs (>10%Chr) have Δ%GC values near 0, some deviate significantly (Δ%GC ≤ -5). We hypothesize that these ERs are strain-specific and not domesticated. Such cases include *Cupriavidus*, *Pseudonocardia*, *Pseudomonas*, *Ruminococcus*, and *Treponema* (Fig. 1b – Betaproteobacteria for *Cupriavidus*; Supplementary Fig. 2 for others). Additionally, our analysis revealed lineage-specific evolutionary groups, including a distinct set of small ERs in Morganellaceae with unusually high Δ%GC values (Supplementary Fig.2 – Morganellaceae).

### Conjugation systems are more prevalent in small ERs than in large ERs

Building on these results, we analyzed the mobility of ERs, focusing on Alphaproteobacteria, Betaproteobacteria, and Gammaproteobacteria. Using ConjScan ^32^, we assessed the conjugative capabilities of ERs and found that small ERs were significantly more likely to encode conjugation systems compared to large ERs, which rarely exhibited such systems (Supplementary Fig. 4, Supplementary Data 3). These findings provide further evidence for distinct evolutionary trajectories of small and large ERs, with the latter showing a higher degree of domestication. To summarize the overall prevalence and distribution of ERs, we found that approximately 50% of bacterial genomes contain ERs. Small and large ERs exhibit distinct distributions across bacterial taxa (Fig. 2, Supplementary Data 4). ER-rich families span diverse bacterial lineages, while large ERs are concentrated within specific lineages, particularly Proteobacteria. This analysis underscores how bacterial lineages have evolved distinct genome architectures, ranging from genomes without ERs to those with small, often transient ERs, to lineages harboring conserved large ERs. All processed data are compiled within Supplementary Data 1, 2, 3 and 4, providing an open resource to explore ER diversity and domestication patterns across bacterial taxa.

**Figure 2:**
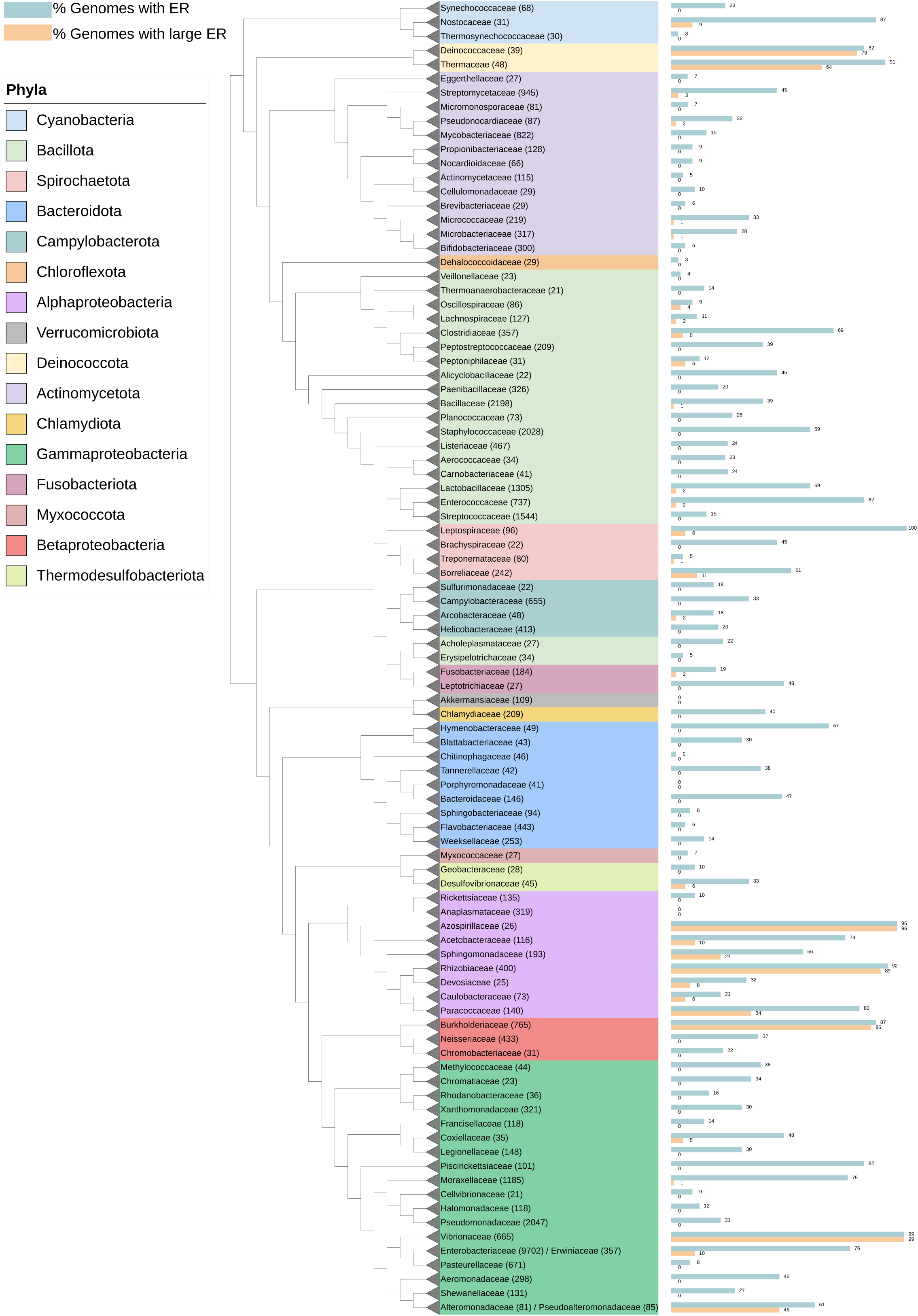
Mapping of ER and large ER onto a reference tree of bacteria. The bacterial phylogeny is based on the GTDB classification ^46^. A subtree was extracted from the GTDB phylogeny, containing 32,900 leaves corresponding to the families of interest (see Supplementary Data 6). For clarity, all leaves belonging to the same family were collapsed.

### Replication termination synchrony with the main chromosome is a common feature of large ERs

We conducted marker frequency analysis (MFA) on 13 bacterial species containing large ERs across five taxonomic classes: Alphaproteobacteria, Betaproteobacteria, Gammaproteobacteria, Deinococci, and Spirochaetia. These strains encompassed a diverse range of large ER sizes and evolutionary origins, representing genera such as *Allorhizobium*, *Sinorhizobium*, *Brucella*, *Paracoccus*, *Cereibacter*, *Burkholderia*, *Cupriavidus*, *Pseudoalteromonas*, *Deinococcus* and *Leptospira*.

MFA was performed during both exponential and stationary growth phases to determine relative copy numbers and observe dynamic replication timing across replicons. During the stationary growth phase, large ERs consistently exhibited the same copy number as the chromosomes (Supplementary Fig. 5), suggesting the presence of a regulatory mechanism that ensures equal copy numbers between chromosomes and large ERs. Using this 1:1 ratio in the stationary phase, we analyzed data from the exponential growth phase to infer the relative replication timing of large ERs, as previously demonstrated for *V. cholerae* ^21^. In the exponential phase (Fig. 3), most replicons displayed “roof” profiles, characterized by peaks at the predicted origins (*ori*) and troughs at the termini (*ter*), indicative of bidirectional replication from *ori* to *ter*. Certain replicons, such as those in *Allorhizobium ampelinum*, *Paracoccus aminophilus*, and *Pseudoalteromonas translucida*, did not exhibit the “roof” shape but instead displayed a unique slope, suggesting unidirectional replication. Notably, the slopes across all replicons in each strain were uniform, indicating that replication speed is identical between the chromosome and large ERs ^33^, likely due to the shared use of the nucleotide pool and replication machinery.

**Figure 3:**
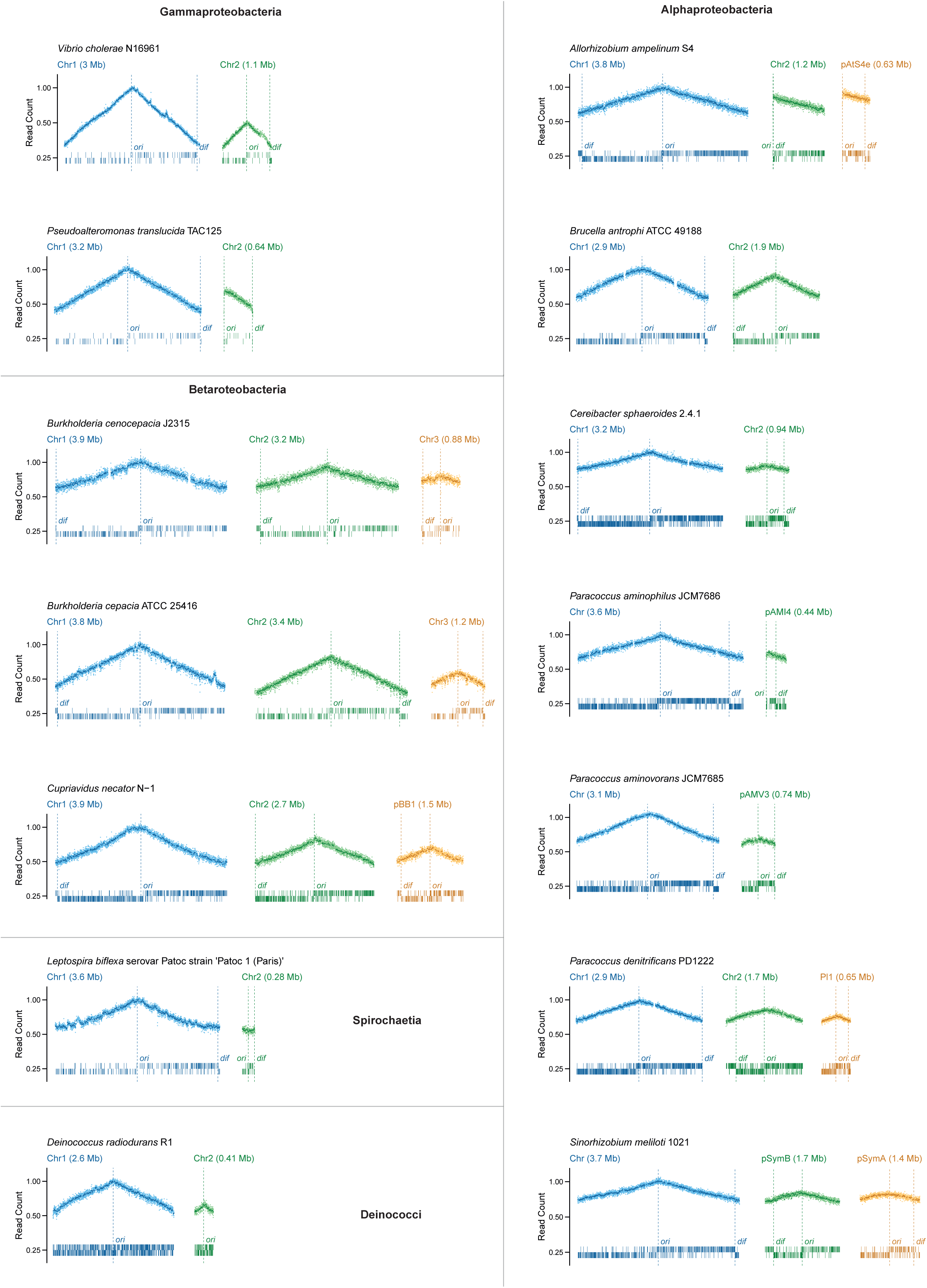
Marker frequency analysis of 13 strains containing large ERs. MFA was performed on gDNA extracted from bacteria in the exponential growth phase. The main chromosome is shown in blue, while ERs are depicted in green and yellow. Read counts were normalized to the *ori* of the chromosome. Darker points represent normalized read counts per 10 kb windows, lighter points correspond to normalized read counts per 1 kb windows. The positions of *ori* and *dif* sites are indicated. Sticks represent the polar distribution of KOPS on the top and bottom strands of the respective replicons.

Across all strains, the main chromosome consistently exhibited the highest initial read counts at *ori*, indicating that it initiates replication before all large ERs. Remarkably, replication termination timing across replicons tended to synchronize. This is evident on the plots (Fig. 3), where the read coverage at the *ter* is equal across replicons, demonstrating that replication ends with similar timing for chromosomes and large ERs. While slight deviations were observed (e.g. the third replicons of *Burkholderia* strains and *Allorhizobium* terminated replication slightly earlier than the main chromosome), these ERs still showed termination timing more aligned with the chromosome than with initiation synchrony.

Our findings show that large ERs across diverse bacterial taxa tend to synchronize their replication termination with the main chromosome, regardless of their evolutionary origin or replication mode (bidirectional or unidirectional). This tendency, combined with the uniform replication speed across replicons and a consistent 1:1 chromosome-to-ER copy number ratio, suggests that coordinated replication dynamics are likely critical for maintaining genome stability in multipartite bacterial genomes, even when large ERs replicate unidirectionally.

### Larges ERs interacts with the main chromosome through origin of replication and along their replichores

To analyze inter-replicon interactions in multipartite bacterial genomes, we performed Hi-C experiments on exponentially growing cultures of five Alphaproteobacteria, three Betaproteobacteria, and one Spirochaetes to explore the interactions between replicons in different multipartite genomes (Fig. 4, Supplementary Fig. 6). As expected, and already observed for *V. cholerae* ^21^, and *A. fabrum* ^34^, the different large replicons in the nine resulting contact maps were characterized as well-individualized entities, each exhibiting a strong diagonal signal reflecting frequent contacts between adjacent loci. The larger ones presented local domains of increased contact frequencies, separated by barriers, as already observed for all bacteria investigated so far ^35^. All large replicons also exhibited a more or less pronounced signal in the opposite diagonal that reflects the bridging of left and right replichores by cohesin complexes in other bacteria ^35^.

**Figure 4:**
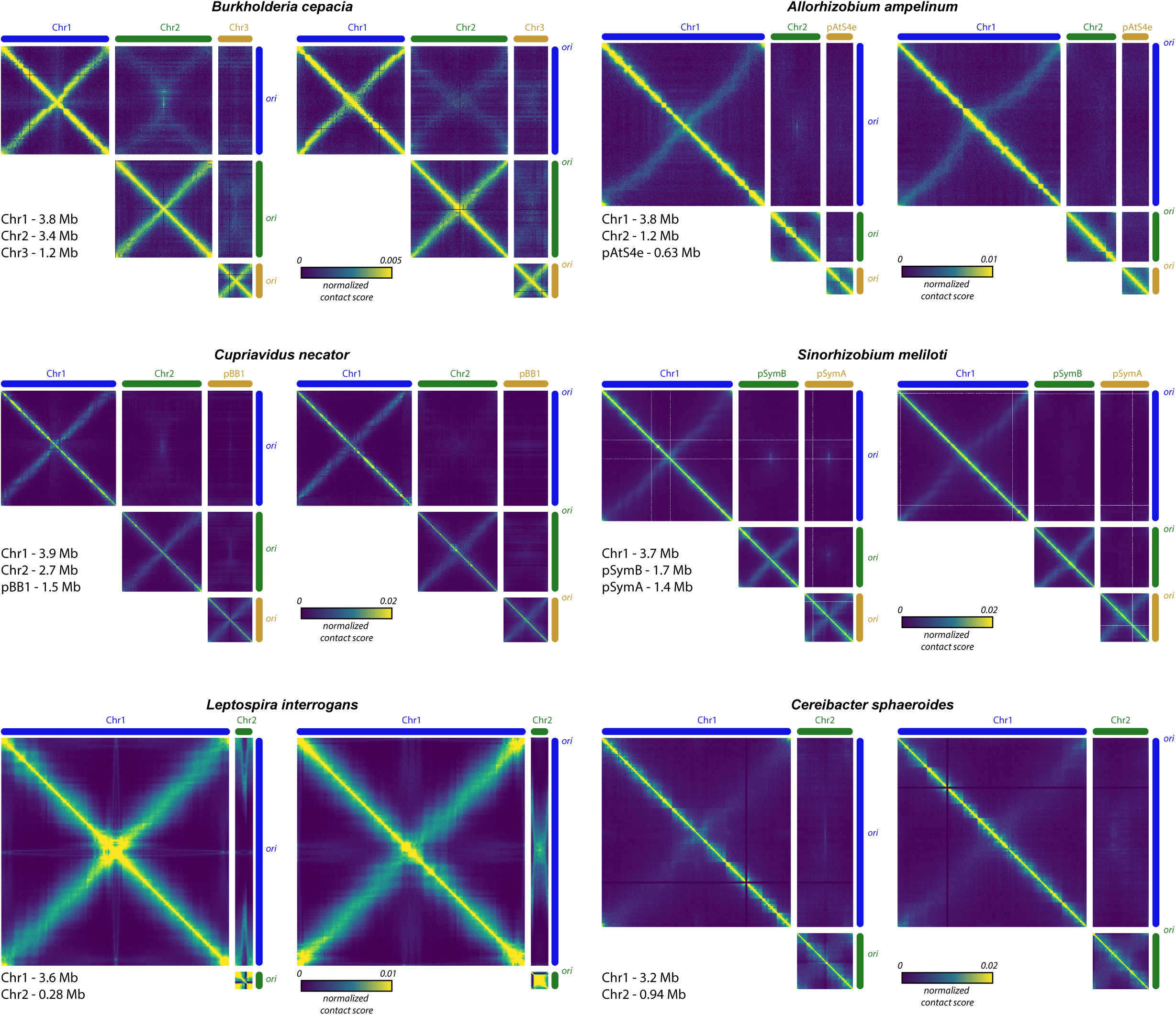
Hi-C interaction matrices between chromosome and large ERs. Normalized Hi-C contact maps of six exponentially growing multipartite bacterial strains. For each strain, two sets of matrices are shown, each with a different replicon orientation. In the left panel, *ori* regions are centered to highlight *ori*-*ori* interactions in the inter-replicon matrices. In the right panel, *ori* regions are positioned at the extremities to emphasize trans-interactions between replichores and potential interactions with *ter* regions in the inter-replicon matrices. For each set of matrices, the replicons are oriented the same way. The x- and y-axes represent the replicons, with *ori* positions marked along the axes. The color scale indicates the frequency of contacts between genomic regions (arbitrary units), ranging from dark blue (rare contacts) to yellow (frequent contacts).

Interestingly, large ERs also exhibit trans-interactions with other replicons. In most species analyzed, we detected similar *ori*-*ori* contacts between chromosome and ERs, including many RepABC replicons (as in *A. fabrum* ^25,34^) as well as replicons with other types of replication systems (Fig. 4). These interactions are strongly represented in our analysis and appear to be specific to the *ori* regions of the interacting replicons. It is important to note that *ori*-*ori* interactions are not exclusive to the chromosome and large ERs as numerous small ERs of the analyzed strains also interact with the chromosome through their *ori* (Supplementary Fig. 6). In some genomes, such as *C. necator*, *A. ampelinum*, *C. sphaeroides*, and B*. cepacia*, we observe a clear interaction pattern extending from the *ori* of the chromosome and large ERs toward the replication termination regions (Fig. 4). These termination regions were inferred from the MFA (Fig. 3), and the observed trans-interactions follow this *ori*-to-termination pattern. Although weaker than *ori*-*ori* interactions, these interactions span much larger regions of the replicons. An especially striking case is *Leptospira interrogans*, where these interactions are highly pronounced.

## DISCUSSION

### A framework for classifying ERs into two groups: small and large ERs

Our analysis demonstrates that two easily recognizable sequence features, Δ%GC and %Chr, are effective in distinguishing different groups of ERs. A general threshold between small and large ERs is observed around 10% of %Chr, although this limit is not absolute. Small ERs exhibit a wide range of Δ%GC values, with their absolute Δ%GC decreasing as ER size increases. Notably, the Δ%GC peak is typically located slightly below 0 in most taxonomic groups. In contrast, large ERs, particularly prominent in Alphaproteobacteria, Betaproteobacteria, and Gammaproteobacteria, tend to have Δ%GC values closer to 0 and can approach sizes comparable to the chromosome. These two groups of ERs overlap in our plots, yet the shift in the central point of Δ%GC distribution suggests a significant difference in their relationship to their host cells.

This difference is further underscored by the presence of conjugative systems, which are more common in small ERs than in large ERs. Large ERs appear to be more evolutionarily integrated with their host genomes. Additionally, some outliers exist among the large ERs, characterized by %Chr greater than 10% and Δ%GC values further from 0, possibly representing distinct evolutionary lineages or more recently acquired elements, such as large conjugative replicons in Alphaproteobacteria.

### Conserved principles of ER domestication in multipartite genomes

Our MFA and Hi-C data reveal consistent features of multipartite bacterial genomes, despite their independent evolution across diverse lineages. One common observation is the presence of *ori*-*ori* interactions between the main chromosome and both large and small ERs. These interactions align with similar findings in systems like *A. tumefaciens* and *Brucella* species, where they play a role in coordinating genome segregation ^25,34,36,37^. Our results suggest that *ori*-*ori* interactions represent a broadly conserved strategy that has independently evolved across bacterial lineages.

A key finding is the synchronized replication termination between the chromosome and large ERs, along with weak interactions spanning the replichores of these replicons. This synchronization likely reflects the need for large replicons to terminate replication simultaneously and at the same cellular location, the closing septum. At this site, mechanisms such as FtsK-mediated DNA translocation and XerCD-mediated recombination ensure efficient chromosome segregation before cell division. FtsK, a DNA translocase, directs chromosome segregation by recognizing specific sequence motifs called KOPS (FtsK Orienting Polar Sequences, 5’-GGGNAGGG-3’), which guide its movement toward the *ter* ^38,39^. In the *ter* region, XerCD recombinases act on *dif* sites to resolve chromosome dimers ^40^. Large ERs exhibit polarized KOPS sequences extending from *ori* to a *dif* site in the *ter* region (Fig. 3), supporting the idea that FtsK operates on these replicons in a chromosome-like manner. This mechanism is critical for the segregation of large ERs and its importance has been demonstrated in *Enterobacteriaceae* and *V. cholerae* ^41,42^. An exception is *D. radiodurans*, where KOPS are abundant but not polarized (Fig. 3) likely due to its high GC content (>65%).

Another important finding is the uniform replication fork speed observed across chromosomes and large ERs (Fig. 3). This uniformity, together with the inter-replicon interactions along their replichores (Fig. 4), suggests a potential coupling of their replication processes. Notably, this link between replication and inter-replicon interactions is particularly evident in cases where large ERs exhibit atypical replication patterns. For instance, in *A. ampelinum*, Chr2 follows a unidirectional replication mode, while in *C. sphaeroides*, Chr2 termination site is not directly opposed to its origin (Fig. 3). These cases allow us to observe that inter-replicon interactions align with the progression of replication from origin to termination (Fig. 4), reinforcing the idea that replication dynamics shape these interactions. A similar mechanism has been proposed for *V. cholerae*, where inter-replicon replichore interactions initiate between *crtS* and *ori2* and terminate at both *ter* regions, mirroring the concurrent replication of Chr1 and Chr2 ^21^. This coupling could be driven by shared molecular elements, such as helicases and polymerases, as most ERs lack a complete replication system beyond the initiator. Coupled replication may thus provide a robust mechanism to coordinate replication and segregation of multiple replicons.

### Evolutionary perspective

Our findings shed light on the evolutionary trajectory of ERs, from small plasmids to large secondary chromosomes. Upon entering a new host, ERs may evolve through parasitic or mutualistic pathways depending on selective pressures. In the mutualistic pathway, referred to as ER domestication, ERs integrate into the host cell cycle by leveraging existing replication and segregation mechanisms designed for the chromosome.

In the early stages of domestication, ERs establish *ori*-*ori* contacts with the chromosome. This mechanism is sufficient for the segregation of smaller ERs, as evidenced by *ori*-*ori* interactions observed across both small and large ERs in diverse bacterial lineages. However, as ERs increase in size, *ori*-*ori* interactions alone may be inadequate to ensure stable inheritance. To overcome this limitation, large ERs may acquire additional features, such as synchronizing their replication termination with the chromosome and incorporating polarized KOPS sequences. These adaptations enable large ERs to integrate into the host’s Ter-domain segregation system and utilize FtsK/XerCD machinery for efficient segregation during cell division. Further adaptations may enhance the stability of large ERs. For example, *V. cholerae* Chr2 mimics chromosomal organization through the acquisition of *matS* sequences in its *ter* region, which define a Ter macrodomain ^30^, and SlmA-binding motifs, which prevent premature division site formation ^43^.

These features highlight the evolutionary flexibility of large ERs, allowing them to integrate into host systems while adapting to chromosomal mechanisms. This evolutionary trajectory may also explain why ERs never exceed the size of the main chromosome. The bacterial cell cycle imposes strict temporal and spatial constraints that are optimized for the replication and segregation of the chromosome. These limitations inherently restrict the size of large ERs, ensuring they remain smaller than the main chromosome while maintaining stable inheritance.

### Avenues for future research

This study lays the groundwork for exploring new questions in the coordination of multipartite bacterial genomes and ER domestication. The comprehensive dataset and findings presented here open several avenues for further investigation:

- <u>Identifying molecular timers:</u> As discovered in *V. cholerae* with the *crtS* checkpoint ^21^ or in *A. tumefaciens* with CcrM DNA methylation ^28^, it would be valuable to search for mechanisms in other bacterial lineages that synchronize the replication timing of large ERs with the chromosome.
- <u>Examining replication concurrency and coupling:</u> The consistent replication speeds observed between chromosomes and large ERs raise the question of whether their replication processes are concurrent and physically coupled. Testing this hypothesis could reveal shared replication machinery or novel interactions that coordinate multiple replicons within a single cell cycle.
- <u>Investigating domestication intermediates</u>: Some ERs in our dataset display GC content significantly different from their host chromosome, suggesting they are not yet fully domesticated. These replicons represent potential intermediates of domestication and could provide unique insights into the evolutionary steps leading to integration into host genome maintenance systems.

## METHODS

### Genome Dataset

A comprehensive bacterial genome dataset was obtained from the NCBI RefSeq repository, specifically including all complete genomes available as of October 10, 2024. Genomes classified as atypical, metagenome-assembled genomes (MAGs), or originating from large multi-isolate projects were excluded to ensure high-quality genome representation.

To ensure data quality, several filtration steps were applied. Genomes containing replicon names labeled as “shotgun” or “partial” were removed, as these are likely incomplete assemblies, resulting in the exclusion of 60 genomes. Additionally, genomes harboring more than 30 replicons were excluded (2 genomes), a number considered implausible for truly complete bacterial genomes. Additionally, we filtered out genomes with any replicon smaller than 1 kb (55 genomes), as these may represent sequencing artifacts or incomplete assemblies. Following these cleaning steps, a final dataset of 43,121 genomes was retained for analysis (Supplementary Data 1).

### Extrachromosomal Replicon Analysis

Chromosomes were defined as the largest replicons within each genome, while all other replicons (i.e. plasmid, megaplasmid, chromid, secondary chromosome…) were defined as extrachromosomal replicons (ERs). Two key features of extrachromosomal replicons (ERs) were calculated: the difference in GC content between each ER and its corresponding chromosome (Δ%GC) and the relative size of each ER compared to the chromosome (%chr). ERs were then plotted based on their Δ%GC and %Chr values at various taxonomic levels, including class, order, and family. Only taxa with more than 100 ERs available in our dataset were included in these visualizations, and the plots are provided in the Supplementary Fig. 3. To analyze the prevalence of ERs across various bacterial taxa, we calculated the percentage of genomes that contained no ERs, those with ERs, and those with large ERs (defined as ERs with %Chr greater than 10%). Additionally, we determined the overall prevalence of ERs, along with the proportion of small ERs (%Chr < 10%) and large ERs (%Chr ≥ 10%) at different taxonomic levels (class, order, family, and genus). These values were calculated by dividing the number of ERs in each size category by the total number of genomes within each taxonomic group. The results of these analyses are presented in Supplementary Data 1

#### Mobility of ERs in Alpha-, Beta-, and Gammaproteobacteria

Mobility was assessed using MacSyFinder v2.1.3 with the CONJScan model ^44,45^. For ERs classified as conjugative (pCONJ) or decayed conjugative (pdCONJ), the conjugation system type was indicated. ERs containing only a relaxase were categorized as mobilizable (pMOB), while ERs lacking mobility-associated genes were designated as non-mobile (pMOBless). The results of these analyses are presented in Supplementary Data 3. The R code used for these analyses are provided in Supplementary Data 5.

#### Mapping of ER and large ER on a reference tree of bacteria

The phylogeny of bacteria is based on the phylogeny provided by GTDB ^46^. The GTDB phylogeny was used to extract a subtree containing 32900 leaves belonging to the families of interest (see Supplementary Data 6). To represent bacterial diversity, 95 families were selected for analysis.

### Strains and growth conditions

The bacterial strains tested in this study were *Allorhizobium vitis* S4 (formerly *Agrobacterium vitis*), *Brucella anthropi* ATCC 49188 (formerly *Ochrobactrum anthropi*), *Burkholderia cenocepacia* J2315, *Burkholderia cepacia* ATCC 25416, *Cereibacter sphaeroides* 2.4.1 (formerly *Rhodobacter sphaeroides*), *Cupriavidus necator* N-1, *Deinococcus radiodurans* R1, *Leptospira biflexa* serovar Patoc strain ‘Patoc 1 (Paris)’, *Leptospira interrogans serovar Manilae strain L495, Paracoccus aminophilus* JCM 7686, *Paracoccus aminovorans* JCM 7685, *Paracoccus denitrificans* PD1222, *Pseudoalteromonas translucida* TAC125 (formerly *Pseudoalteromonas haloplanktis*), and *Sinorhizobium meliloti* 1021.

All strains, except *A. vitis*, *P. translucida*, *D. radiodurans*, *L. biflexa*, and *L. interrogans,* were grown in LB Lennox medium at 30°C. *A. vitis* was grown in TY medium (bacto-tryptone 5 g/L, yeast extract 3 g/L, CaCl₂·2H₂O 0.87 g/L) at 30°C, *P. translucida* in TYP medium (bacto-tryptone 16 g/L, yeast extract 16 g/L, NaCl 20 g/L) at 20°C, *D. radiodurans* in TSB medium (Tryptic Soy Broth) at 30°C, and *Leptospira* spp. in EMJH medium at 30°C.

### Marker Frequency Analysis

#### Preparation of bacterial samples in exponential and stationary phases

Overnight cultures were initiated from fresh streaks on solid medium by inoculating 3 mL of liquid medium and incubating for 24 hours with shaking (150 rpm). To obtain exponential phase cultures, 0.1 mL of the overnight culture was transferred into 100 mL of fresh medium and incubated in a shaking water bath (180 rpm) until reaching an OD₄₅₀ of 0.2–0.4, except for *L. biflexa*, which was grown to an OD₄₂₀ of 0.2. For exponential phase samples, bacteria were harvested by centrifuging the total culture volume in two 50 mL Falcon tubes at 4°C for 10 minutes at 8000 rpm. The supernatant was discarded, and the bacterial pellets were immediately stored at -20°C to halt replication.

For stationary phase samples, incubation of overnight cultures was continued for an additional 8 hours, except for *L. biflexa*, where cultures were grown until an OD₄₂₀ of 0.6. Cultures (0.5 mL, or 50 mL for *L. biflexa*), were centrifuged in 1.5 mL Eppendorf tubes at room temperature for 1 minute at 13,000 rpm. The supernatant was removed, and the bacterial pellets were stored at -20°C.

#### Genomic DNA extraction and sequencing

Genomic DNA was extracted from bacterial pellets obtained from both exponential and stationary phases. For all strains except *P. translucida* TAC125, the GeneMATRIX Bacterial & Yeast Genomic DNA Purification Kit (EurX) was used. DNA extraction for *P. translucida* was performed using the DNeasy Blood & Tissue Kit (Qiagen). The extracted genomic DNA was quantified using a Qubit Fluorometer (Life Technologies), yielding DNA concentrations ranging from 30 to 300 ng/μL.

DNA Sequencing. Sequencing libraries were prepared according to the Truseq DNA PCR-Free library preparation protocol (Illumina). Equimolar pools of libraries were sequenced using a Hiseq 2500 system (paired-end 100 bp) or Nextseq 500 system (pair-end 150 bp) (Illumina), to obtain 15-20M reads per sample.

Sequence analysis. Sequence reads were mapped to the reference sequences with Bowtie2 with default settings ^47^ and the number of reads per 1 kb window were extracted from the output with a self-made Perl script. Then, a correction factor for each 1 kb window was calculated from the stationary-phase reads and was used to normalize the exponential-phase reads (as on Fig. S15, panels A and B in ^21^). Windows covering repeated DNA sequences were removed using the R2R scripts ^19^ and manual curation. The numbers of reads per 1 kb window were then scaled to the ori of the main chromosome and plotted with a logarithmic ordinate ^19^.

MFA was used to observe replication patterns in replicons larger than 250 kb, which served as the cutoff for our analysis. In replicating bacteria (exponential phase), MFA graphs show the highest number of reads at the origin of replication and the lowest at the terminus, indicating ongoing DNA replication. Conversely, non-replicating bacteria (stationary phase) exhibit flat MFA graphs, reflecting a lack of active replication.

Stationary phase sequencing data served two purposes: (1) to normalize the exponential phase sequencing results and (2) to determine the relative copy numbers of the chromosome and secondary replicons. We only included replicons with copy numbers similar to the main chromosome in the stationary phase to make accurate conclusions about their relative replication timing during the exponential phase. For two bacterial strains, *L. biflexa* and *D. radiodurans*, the stationary phase MFA plots did not show the expected flat shape. Despite multiple attempts to cultivate them for extended periods to reach the stationary phase, we consistently observed non-flat profiles (Supplementary Fig. 5), likely due to specific growth characteristics of these organisms. Nevertheless, we present their data, as the exponential phase results strongly suggest replication synchrony between the main chromosome and their large extrachromosomal replicons.

### Hi-C

The different bacterial strains were grown until mid-exponential phase (OD = 0.3) in 80 mL of LB. Bacteria were then treated with formaldehyde at a final concentration of 3%. Cultures were then incubated for 30 min under shaking at RT followed by incubation at 4°C during another 30 min. Formaldehyde was quenched by adding three times the volume of Glycine 2.5 M followed by an incubation of 20 min under shaking. Pellets were recovered by a centrifugation of 10 min at 10,000 x g and at 4°C, washed in PBS and re-centrifuge using the same settings. Pellets were then frozen in dry ice and stored at -80°C until use. Hi-C libraries were generated using the ARIMA kit (Arima Genome-Wide Hi-C Kit). Samples were first resuspended in 1 mL of sterile water and transferred to 2 mL precellys tubes containing glass beads of 0.1- and 0.5-mm diameter (Precellys – Bertin Technology). Cells were disrupted using the precellys apparatus (Precellys Evolution) and the following program (7500 rpm, 6 cycles 30 sec ON / 30 sec OFF, 4°C). Tubes were then centrifuged for 1 min at 1000 g and 700 µL of lysate was recovered and transferred to a new 1.5 mL eppendorf tube. Tube was centrifuged for 20 min at 16,000 g, 4°C. Supernatant was carefully removed and pellet was resuspended in 45 µL of water. We then followed the ARIMA protocol and the final library was purified using AMPure beads and eluted in 130 µL of water before processing for sequencing. HiC genomic libraries were first sheared at a mean size of 500 bp using a covaris S220 apparatus, processed on streptavidin beads using the colibri kit (ThermoFisher) as previously described ^48,49^. Libraries were sequenced on NextSeq apparatus (2 x 35 bp). Contact maps were generated using Hicstuff (bowtie2 - very sensitive local mode – mapping quality of 30) and the different reference genomes. Contact maps were then binned at different resolutions (1 kb, 2 kb and 5 kb), balanced and displayed using cooler and different R packages ^50^.

## Data Availability

The sequencing data generated in this study have been deposited in the European Nucleotide Archive under project accession code PRJEB85805.

## Acknowledgments

We thank the following individuals for providing bacterial strains and guidance on cultivation methods: Ludovic Vial, Evelyne Krin, Franck Pasta, Denis Faure, Pamela Brown, the Collection de l’Institut Pasteur (CIP). Our gratitude extends to Marc Monot and Juliana Pipoli Da Fonseca (Biomics core facility, IP, C2RT) for their technical support in sequencing. We also thank the PF Milieux and PF Matériel teams for daily assistance. We acknowledge the LABGeM (CEA/Genoscope & CNRS UMR8030), France Génomique and French Bioinformatics Institute national infrastructures (funded through Investissement d’Avenir and France 2030 programs managed by Agence Nationale pour la Recherche, contracts ANR-10-INBS-09, ANR-11-INBS-0013 and ANR-21-ESRE-0048) for their support via the MicroScope annotation platform. This work was supported by the Institut Pasteur; Institut National de la Santé et de la Recherche Médicale (INSERM); Centre National de la Recherche Scientifique [CNRS-UMR 3525]; French National Research Agency [ANR-19CE12-0001]; [ANR-23-CE12-0005]; [ANR-10-LABX62-IBEID] ; Fondation pour la Recherche Médicale [Equipe FRM EQU202103012569] and FDM202106013531, to M.L. J.C. was supported by [ANR-10-LABX-62IBEID], [ANR-19-CE12-0001], the Institut Pasteur Roux-Cantarini fellowship. EK was funded by National Science Centre, Poland [2023/49/N/NZ2/01061].

## Competing interests statement

The authors declare no competing interests.

---

**Supplementary Figure 1 : Distribution of chromosomes and ERs by phylum based on GC Content and size (Logarithmic Scale)**

Chromosomes (black) and extrachromosomal replicons (ERs; purple) from the five phyla with the highest number of ERs in the RefSeq database, along with data from other phyla, are analyzed. Replicons are plotted by size (bp) and GC content (%), using logarithmic scale to enhance the resolution of both large and small replicons. The total numbers of chromosomes (N(chr)) and ERs (N(ER)) are indicated on each plot. The number of chromosomes corresponds to the number of genomes analyzed per plot, as chromosomes were defined as the largest replicons in each genome, while ERs were categorized as all other replicons within the same genome.

**Supplementary Figure 2 : Comparison of different methods for ER classification based on size and GC content** Size is expressed in base pairs (bp), as a percentage of the chromosome size (%Chromosome), and as a percentage of the whole genome size (%Genome). GC content is expressed either as %GC or as Δ%GC (the difference between the GC content of the ER and the chromosome from the same genome). Only ER replicons are plotted, colored according to their classification in the RefSeq database: secondary and tertiary chromosomes (blue), megaplasmids (red), plasmids (orange), and undetermined (gray).

**Supplementary Figure 3 : Classification of ERs by size and GC content across families with more than 100 ERs** ERs from bacterial families with more than 100 ERs in our dataset are plotted based on their size, expressed as a percentage of the chromosome (%Chromosome), and the difference in GC content compared to the chromosome of the same cell (Δ%GC). For each family, ERs from up to five genera with the highest number of large ERs (defined as those exceeding 10% of the chromosome size) are highlighted. Dashed lines indicate the 10% Chr threshold, which serves as our proposed boundary between small and large ERs, and Δ%GC = 0, where ERs share the same GC content as their corresponding chromosome.

**Supplementary Figure 4 : Mobility of small and large ERs.**

Bar plot showing the percentage of ERs classified as conjugative (pCONJ), decayed conjugative (pdCONJ), mobilizable (pMOB), or non-mobile (pMOBless) in Alpha-, Beta-, and Gammaproteobacteria. pCONJ (conjugative plasmids) encode all the key genes required for conjugation, i.e., mating pair formation (MFP) machinery and a relaxase. pdCONJ (decayed conjugative plasmids) encode an incomplete MFP machinery and a relaxase. pMOB (mobilizable plasmids) encode only a relaxase, lacking the MFP machinery, meaning they cannot transfer independently but can be mobilized by a co-residing conjugative plasmid. pMOBless (non-mobilizable plasmids) do not encode any conjugation-related machinery. This classification is based on ConjScan analysis ^32^.

**Supplementary Figure 5 : Marker frequency analysis of 13 strains containing large ERs in stationary phase**

MFA was performed on gDNA extracted from bacteria in the stationary growth phase. The main chromosome is shown in blue, while ERs are depicted in green and yellow. Read counts were normalized to the ori of the chromosome. Darker points represent normalized read counts per 10 kb windows, lighter points correspond to normalized read counts per 1 kb windows. The positions of ori and dif sites are indicated. Sticks represent the polar distribution of KOPS on the top and bottom strands of the respective replicons. Note that for two bacterial strains, *L. biflexa* and *D. radiodurans*, the stationary-phase MFA plots did not exhibit the expected flat profile. Despite multiple attempts to cultivate them for extended periods to reach stationary phase, we consistently observed non-flat profiles, likely due to specific growth characteristics of these organisms.

**Supplementary Figure 6 : Hi-C interaction matrices between chromosome and all ERs**

Normalized Hi-C contact maps of nine exponentially growing multipartite bacterial strains. For each strain, ori regions are set to 0 to emphasize trans-interactions between replichores and potential interactions with ter regions in the inter-replicon matrices. In all matrices, the replicons are oriented the same way. The axes represent the replicons, following the same color scheme as in Fig. 4, with small ERs additionally colored in grey. The color scale indicates the frequency of contacts between genomic regions (arbitrary units), ranging from dark blue (rare contacts) to yellow (frequent contacts).

## Notes

### Competing Interest Statement

The authors have declared no competing interest.

